# SARS-CoV-2 targets cortical neurons of 3D human brain organoids and shows neurodegeneration-like effects

**DOI:** 10.1101/2020.05.20.106575

**Authors:** Anand Ramani, Lisa Müller, Philipp Niklas Ostermann, Elke Gabriel, Pranty Abida-Islam, Andreas Müller-Schiffmann, Aruljothi Mariappan, Olivier Goureau, Henning Gruell, Andreas Walker, Marcel Andrée, Sandra Hauka, Torsten Houwaart, Alexander Dilthey, Kai Wohlgemuth, Heymut Omran, Florian Klein, Dagmar Wieczorek, Ortwin Adams, Jörg Timm, Carsten Korth, Heiner Schaal, Jay Gopalakrishnan

## Abstract

COVID-19 pandemic caused by SARS-CoV-2 infection is a public health emergency. COVID-19 typically exhibits respiratory illness. Unexpectedly, emerging clinical reports indicate that neurological symptoms continue to rise, suggesting detrimental effects of SARS-CoV-2 on the central nervous system (CNS). Here, we show that a Düsseldorf isolate of SARS-CoV-2 enters 3D human brain organoids within two days of exposure. Using COVID-19 convalescent serum, we identified that SARS-CoV-2 preferably targets soma of cortical neurons but not neural stem cells, the target cell type of ZIKA virus. Imaging cortical neurons of organoids reveal that SARS-CoV-2 exposure is associated with missorted Tau from axons to soma, hyperphosphorylation, and apparent neuronal death. Surprisingly, SARS-CoV-2 co-localizes specifically with Tau phosphorylated at Threonine-231 in the soma, indicative of early neurodegeneration-like effects. Our studies, therefore, provide initial insights into the impact of SARS-CoV-2 as a neurotropic virus and emphasize that brain organoids could model CNS pathologies of COVID-19.

**One sentence summary:** COVID-19 modeling in human brain organoids

## Introduction

The novel coronavirus disease 2019 (COVID-19) caused by Severe Acute Respiratory Syndrome Coronavirus 2 (SARS-CoV-2) is spreading worldwide, and the outbreak continues to rise, posing a severe emergency (*1*). Understanding the biology of the current COVID-19 pandemic is a high priority for combatting it efficiently. Thus, it is essential to gain initial insights into the infection mechanisms of SARS-CoV-2, including its target cell types and tropism, to contain its short- and long-term effects on human health. Furthermore, it is vital to establish an experimental system that could allow designing measures on how to stop viral replication and protect human health rapidly. However, practical problems associated with the isolation and handling of highly infective viral strains and lack of reliable in vitro human model systems that can efficiently model COVID-19 hamper these efforts.

Clinical symptoms of COVID-19 patients include upper respiratory tract infection with fever, dry cough, and dyspnea, indicating that the respiratory tract is the first target (*2*). However, emerging case reports showed that patients infected with SARS-CoV-2 suffered a sudden and complete loss of the olfactory function, stroke, and other severe neurological symptoms (*3–7*). All of these indicate that SARS-CoV-2 is neurotropic and could infect the central nervous system (CNS) (*8–10*). Importantly, it remains unclear whether SARS-CoV-2 is vertically transmitted to fetuses, where it could impair neurodevelopment (*11*). Since coronaviruses share a similar structure, it is likely that SARS-CoV-2 exhibits the same infection mechanism and possibly invades into the brain (*12*). Indeed, a clinical report detected the presence of viral RNA in autopsy brain samples (*13*). Thus, at this point, it is essential to test whether SARS-CoV-2 infects human neurons and productively replicates in the CNS.

Examining a potential neurotropism of SARS-CoV-2, it is essential to employ a suitable in vitro human model system that can recapitulate the physiological effect of SARS-CoV-2 infection. In this regard, the recently emerged human brain organoids that closely parallel the complex neural epithelium exhibiting a wide diversity of cell types could serve as a suitable model system to test the neurotoxic effects of SARS-CoV-2. Induced Pluripotent Stem Cells (iPSCs)-derived human brain organoids have revealed useful insights into human brain development and helped to model a variety of neurological disorders (*14*) (*15–17*) (*18*) (*19*). Notably, others and our work using brain organoids have revealed unprecedented insights into infection mechanisms, target cell types, and the toxicity effects of the Zika virus (ZIKV) during the recent ZIKV epidemic. These studies validate organoids as a tool for studying not only genetic but also environmental hazards to human brain (*20*) (*21*) (*22*).

Here, we report that SARS-CoV-2 readily targets cortical neurons but not neural stem cells of 3D human brain organoids. Neurons invaded with SARS-CoV-2 at the cortical area display Tau missorting, Tau hyperphosphorylation and apparent neuronal death. Moreover, we show that although SARS-CoV-2 can readily target brain organoids, SARS-CoV-2 does not productively replicate, suggesting that the CNS may not support the replication of SARS-CoV-2.

## Results

### Isolation of an infectious SARS-CoV-2 virus

We isolated SARS-CoV-2 from an infected patient admitted to our university hospital, University of Düsseldorf (see method section for culturing and propagation). To investigate whether SARS-CoV-2 replicates in inoculated African green monkey kidney cells (Vero CCL-81), we performed realtime quantitative polymerase chain reaction (qPCR) analysis with cell culture supernatant. As expected, the amount of SARS-CoV-2 RNA drastically increased from 0-dpi until 3-dpi **(Figure S1A).** Next, we analyzed the infectivity of generated SARS-CoV-2 particles by propagating virus-containing supernatant to new Vero cells. We confirmed the infection of new Vero cells by the emergence of virus-induced cytopathic effects (CPEs) and an increase in SARS-CoV-2 RNA over 4-dpi. Taken together, we demonstrated the successful generations of a novel SARS-CoV-2 isolate, SARS-CoV-2 NRW-42, derived from a nasopharyngeal and oropharyngeal swab specimen. The sequence access (number PRJNA627229) showed only eight nucleotide exchanges compared to SARS-CoV-2 Wuhan-Hu-1 isolate. To test for potential adaption to the used Vero CCL-81, we whole-genome sequenced virus samples collected from the clinical specimen used for inoculation and virus samples from the supernatant after four days of propagation. Sequence analysis revealed no nucleotide substitution after isolation and propagation in Vero cells. In summary, we isolated an infectious SARS-CoV-2 isolate (NRW-42) with high sequence similarity to Wuhan-Hu-1.

### Isolation and validation of COVID-19 convalescent serum to detect SARS-CoV-2 infection

First, we set out to identify antibodies that can specifically determine SARS-CoV-2 infection. We isolated COVID-19 convalescent serum from blood samples of four independent individuals who recently recovered from COVID-19 (AB1, AB2, AB3, and AB4). Testing them in an enzyme-linked immunosorbent assay (ELISA) that used the S1 domain of the spike protein of SARS-CoV-2 as an antigen revealed that except for AB2, the rest of the convalescent serum contained SARS-CoV-2-specific IgG **(Figure S1B)**. We chose to work with AB4 as it displayed the best immunoreactivity to SARS-CoV-2-exposed brain organoid tissues than the other serum. AB4 specifically recognized SARS-CoV-2-infected Vero cells **(Figure 1A)**.

**Figure 1.**
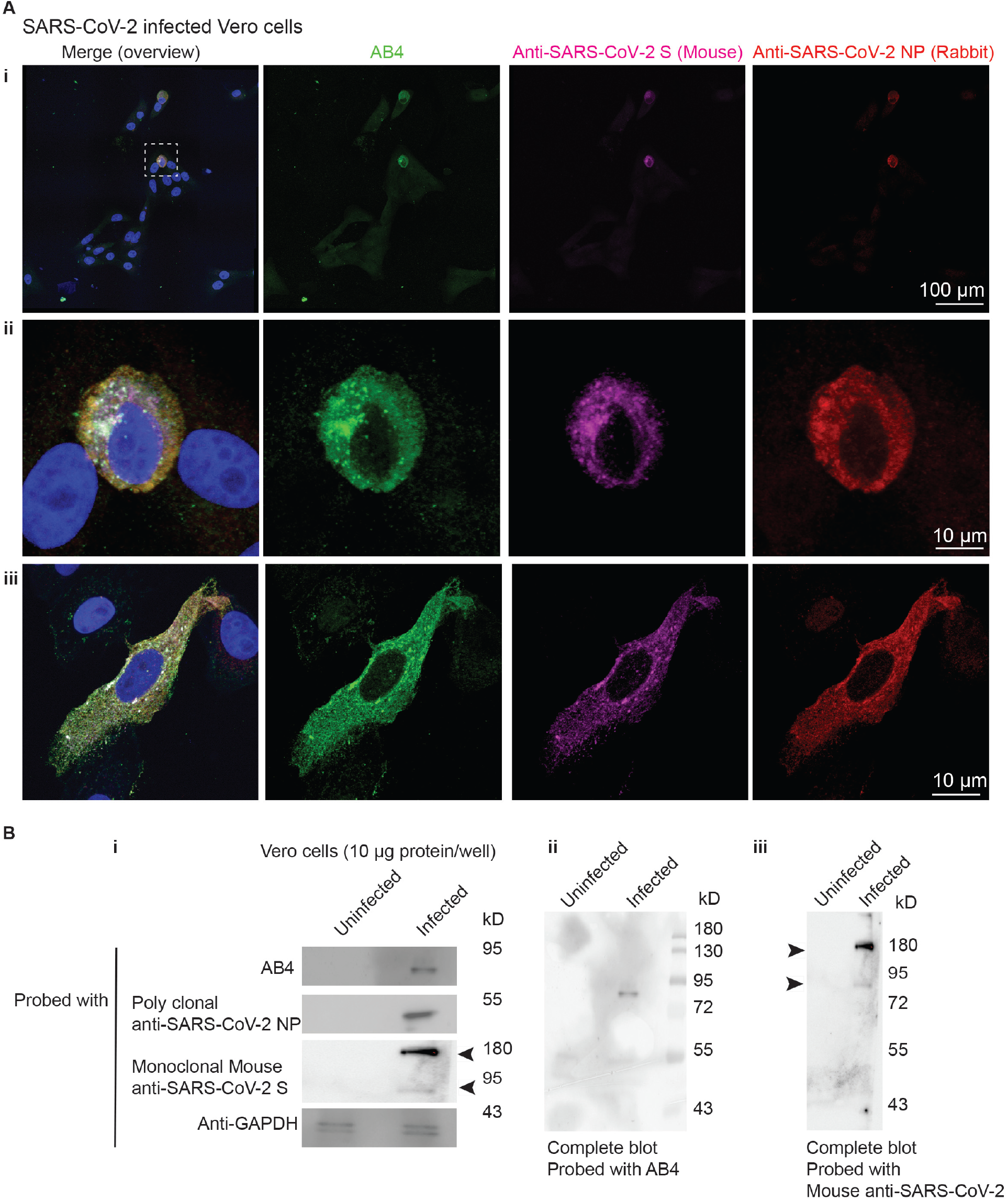
Convalescent serum specifically detects SARS-CoV-2 infected cells. **A.** The convalescent serum AB4 (green) specifically recognizes SARS-CoV-2 infected Vero cells **(i** to **iii).** Mouse monoclonal Anti-SARS-CoV-2 S (magenta) and a polyclonal Anti-SARS-CoV-2 NP (red) validate the specificity of the AB4. At least 60 cells were examined. Figures display scale bars. At least two exemplary imagesof cells **(ii** and **iii)** are shown. **B.** In Western blots, AB4 recognizes a ~75 kD protein band that is similar to the size of the cleaved spike protein in infected Vero cell extracts **(i)**. The same size of the band is also recognized by the monoclonal Anti-SARS-CoV-2 S in addition to ~180 kD corresponding to the sizes of un-cleaved S protein (S0, ~180 kD). Arrowheads mark un-cleaved spike protein (S0, ~180 kD) and the cleaved spike protein (~75 kD). Likewise, polyclonal Anti-SARS-CoV-2 NP recognizes a protein band relevant to nucleoprotein in infected Vero cell extracts. GAPDH blot ensures equivalent protein loading. Representative blots from three independent experiments are given. Whole blots for AB4 **(ii)** and Anti-SARS-CoV-2 S **(iii)** are shown as well.

To further validate the specificity of the AB4, we performed co-immunostaining with a mouse monoclonal Anti-SARS-CoV-2 S and a polyclonal Anti-SARS-CoV-2 NP. These commercial antibodies were raised against the AA 1029-1192 of the S2 domain and AA-1-100 of the nucleoprotein of SARS-CoV-2, respectively. All of these antibodies recognized only SARS-CoV-2-infected Vero cells **(Figure 1A).** In Western blots that used SARS-CoV-2-infected Vero cell extracts, AB4 detected a protein band with an apparent molecular weight of 75 kD, a size similar of the cleaved-spike protein. Both rabbit polyclonal and mouse monoclonal antibodies recognized protein bands around 50 and 180 kDs, sizes similar to the nucleoprotein and uncleaved spike proteins **(Figure 1B)**. Together, these experiments validate that AB4 detects SARS-CoV-2 infection.

### SARS-CoV-2 targets human brain organoids but does not appear to replicate productively

Although the apparent target of SARS-CoV-2 is the respiratory tract, recently reported neurological phenotypes indicate that SARS-CoV-2 has detrimental effects on the CNS and could infect the CNS directly. To test this hypothesis, we generated 3D human brain organoids from two different iPSC lines and exposed them to our SARS-CoV-2 NRW-42 isolate. As described before, organoids exhibit their specific neuronal cell types, which are spatially restricted. For example, apical neural progenitors cells displaying elongated nuclei (NPCs) are aligned to form a lumen that is similar to the ventricular zone (VZ) of the mammalian brain. Cortical neurons are positioned basally to the VZ forming cortical plate **(Figure 2A)** (*23*) (*24, 25*). We exposed at least two different age groups of organoids (Day-15 and Day-60) to SARS-CoV-2 (TCID_50_/mL of 50) and analyzed after 2-and 4-dpi.

**Figure 2.**
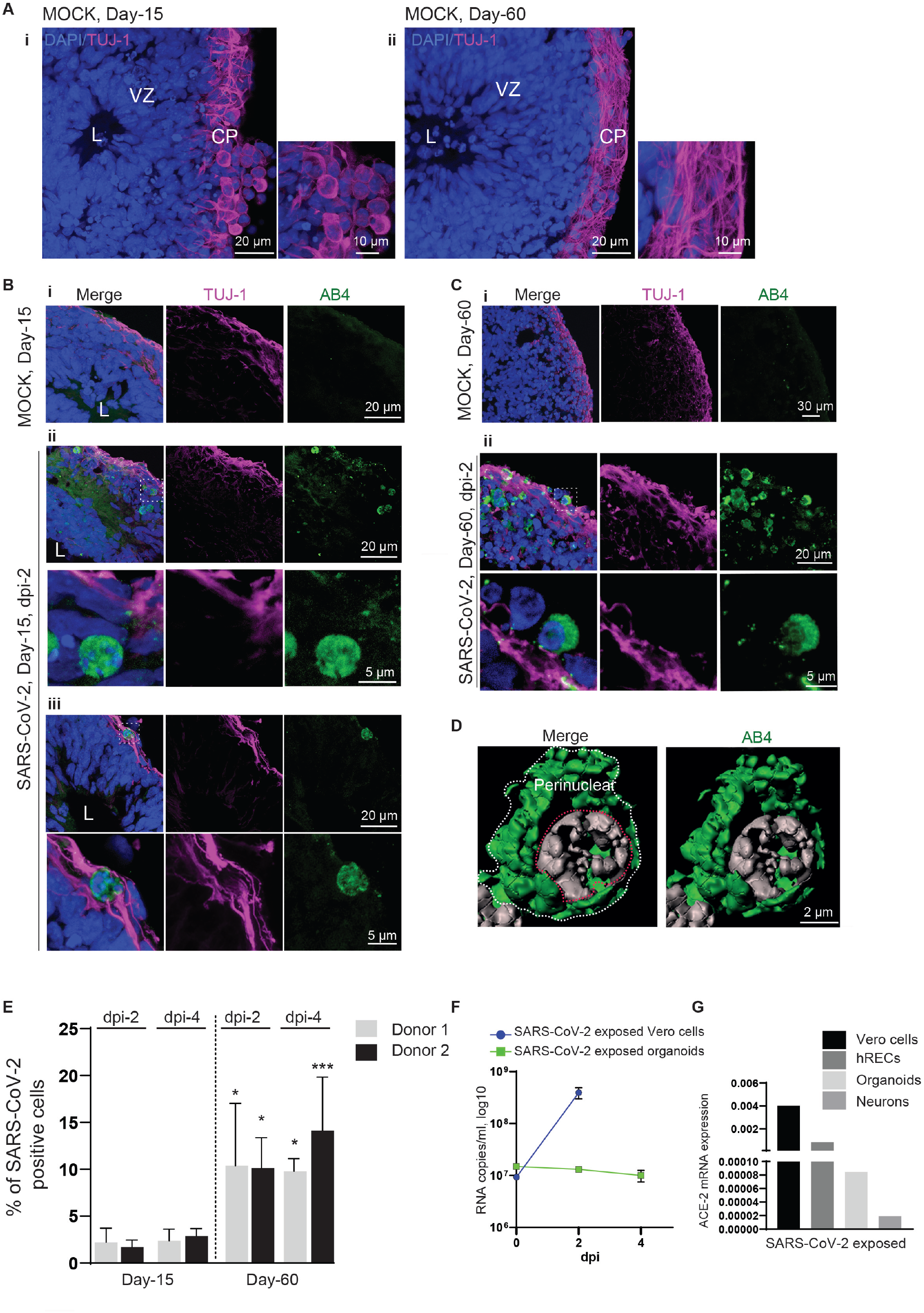
SARS-CoV-2 targets the cortical region of human brain organoids but do not appear to replicate productively. **A.** Mock organoids of two age groups Day-15 **(i)** and-60 **(ii)** display typical cytoarchitecture of brain organoids. L, Lumen, VZ, Ventricular Zone is containing compact and palisade-like elongated nuclei of Neural Progenitor Cells (NPCs, blue) and CP, a cortical plate containing TUJ-1-positive neurons (magenta). Note a distinct difference TUJ-1 labeling pattern between younger (Day-15) and older (Day-60) brain organoid. Figures display scale bars. Representative images from eight organoids cultured in at least three independent batches (n=3) derived from donor-1 (IMR90) iPSC line. **B.** Compared to mock organoids **(i)**, SARS-CoV-2-exposed Day-15 organoids display SARS-CoV-2-positive cells (green) in their outer periphery, a region of the cortical plate **(ii)** that is specified by TUJ-1-positive neurons (magenta). L, the lumen of a VZ, the inner area of an organoid where NPCs are located, is free from SARS-CoV-2-positive cells. Magnified region (dotted while box) is given below. **(iii)** Another example showing the preferential target of SARS-CoV-2 at the cortical plate and not NPCs. Magnified region (dotted while box) is given below. At least eight organoids from four different batches (n=4) are tested. Figures display scale bars. **C.** SARS-CoV-2-exposed Day-60 organoids **(i)**. Compared to Day 15 organoids, Day-60 organoids display an increased number of SARS-CoV-2-positive cells (green) in their cortical plate that is specified by TUJ-1-positive neurons (magenta) **(ii).** Magnified region (dotted while box) is given below, showing the peri-nuclear location of SARS-CoV-2 in cortical neurons. At least eight organoids from four different batches (n=4) are tested. Figures display scale bars. **D.** Subcellular localization SARS-CoV-2 in cortical neurons. High-resolution imaging and de-convolution show peri-nuclear localization of SARS-CoV-2. SARS-CoV-2 (green) and nucleus (grey). Figures display scale bars. Representative images from at least 200 cells examined. **E.** The bar diagram quantifies frequencies of SARS-CoV-2-positive cells in brain organoids derived from two different donor iPSC lines (IMR90 and Crx-iPS, see methods). SARS-CoV-2 shows an enhanced tropism for Day-60 organoids. Note, comparative statistics are shown between different age groups and respective days post infection (dpi) of organoids, and the significance is given as Asterisks in Day-60 groups. There is no significant difference in SARS-CoV-2-positive cells between 2- and 4-dpi within each age groups. At least sixteen organoids from four (n=4) independent batches from each donor were analyzed. One-way ANOVA, followed by Tukey’s multiple comparisons test, *P<0.05, ***P=0.001. Error bars represent mean + SD. **F.** Determination of viral progeny. Supernatants of SARS-CoV-2 exposed Vero cells, and brain organoids were analyzed for viral RNA assessed by qRT-PCR. While an increase in viral RNA was detected in the supernatants of Vero cells, no apparent increase was identified in brain organoid supernatants. Data are obtained from five technical replicates from four (n=4) independent batches of organoids. Error bars represent mean + SD. **G.** Relative fold differences of ACE-2 mRNA expression between Vero cells, human respiratory epithelial cells (hRECs), brain organoids, and iPSC-derived neurons. Data are from multiple technical replicates from two (n=2) independent experiments.

Cryo-sectioning followed by immunofluorescence analysis using convalescent serum indicated that except AB2, all of them could specifically recognize somas of SARS-CoV-2-positive cells in SARS-CoV-2 exposed organoids **(Figure S2A)**. In our opinion, among others, AB4 displayed the best immunoreactivity. Interestingly, AB2, which did not contain S1-reactive IgG, did not label cells in SARS-CoV-2 exposed organoids **(Figure S2Aiv).** A sub-population of cells recognized by AB4 was also labeled by the monoclonal anti-SARS-CoV-2 S antibody, indicating that SARS-CoV-2 could target organoids within 2-dpi **(Figure S2B).**

First, we began analyzing Day-15 organoids, a developmental stage that we used to study ZIKV infections (*20*). At this developmental stage, organoids mostly constitute actively proliferating NPCs at the VZs and a primitive cortical plate containing early neurons **(Figure 2A)**. Testing the target cell types of SARS-CoV-2 in these organoids revealed that SARS-CoV-2 could mostly target the cortical plate specified by pan-neuronal marker TUJ-1 **(Figure 2B-C).** Importantly, the perinuclear localization of SARS-CoV-2 in somas of cortical neurons is similar to the localization pattern of the virus in Vero cells, indicating that SARS-CoV-2 can enter into neuronal cells of brain organoids **(Figure 2D)**.

Of note, although these organoids are abundant with NPCs, we did not detect SARS-CoV-2-positive NPCs at the VZ, suggesting that SARS-CoV-2 preferably targets the cortical region of the brain organoid **(Figure. 2Biii)**. In this respect, the infection route of SARS-CoV-2 is strikingly differing from the recent ZIKV isolate that has a robust neurotropism preferably infecting neural NPCs at the VZ (*20*) (*26*). Analyzing Day-60 organoids revealed that the number of SARS-CoV-2-positive cells was significantly higher than in Day-15 organoids. This suggests that SARS-CoV-2 prefers relatively mature neuronal cell types present in older organoids differentiated from both iPSCs donors **(Figure. 2C and E)**. Since we did not find differences between donors, the rest of the experiments utilized Day-60 organoids generated from donor-2, which displayed signs of cortical maturation as judged by the presence of more MAP2-positive neurons **(Figure S2C).**

Turning our analysis to the later time point of infection (dpi-4) revealed that there was no apparent increase in SARS-CoV-2-positive cells **(Figure 2E).** Corroborating to this, we could not detect an increase in viral RNA in the supernatants between 2- and 4-dpi, indicating that SARS-CoV-2 does not productively replicate in brain organoids until 4-dpi but suggestive of an abortive replication **(Figure 2F).** This is in contrast to recent reports showing SARS-CoV-2 productively infects vascular, kidney, and gut organoids (*27–29*). Notably, angiotensin-converting enzyme 2 (ACE-2), an entry receptor of SARS-CoV-2 is highly expressed in these organoid types. Testing the ACE-2 expression at the level of mRNA via a qRT-PCR revealed that both iPSCs-derived brain organoids and neurons exhibited ~12.5 and 50 folds lesser than human respiratory epithelial cells (hREC), which served as a positive control **(Figure 2G)**.

### SARS-CoV-2-positive neurons reveal aberrant Tau localization and phosphorylation

Next, we identified that the SARS-CoV-2-positive region of the cortical plate is further substantiated by Tau, a microtubule-associated protein that stabilizes neuronal microtubules and promotes axonal outgrowth **(Figure 3)** (*30*). Tau dysfunction is implicated in Alzheimer’s disease and other Tauopathies. Aberrant Tau proteostasis is also a sensitive marker for unspecific lesions of the CNS, such as traumatic brain disorder (*31*). To investigate the consequences of SARS-CoV-2’s effect on cortical neurons, we imaged the Tau localization on SARS-CoV-2-exposed organoids using a pan Tau antibody. Under physiological conditions, Tau is mainly an axonal protein that labels the axons of mature neurons **(Figure. 3Ai-iv and S4A)**. Applying high-resolution imaging followed by de-convolution, we could visualize the localization of Tau exclusively in axons the cortical neurons **(Figure 3Aiv)**. The term Tau missorting is used when Tau protein is mislocalized into a somatodendritic compartment and is observed at the early stages of Tau pathology (*32*).

**Figure 3.**
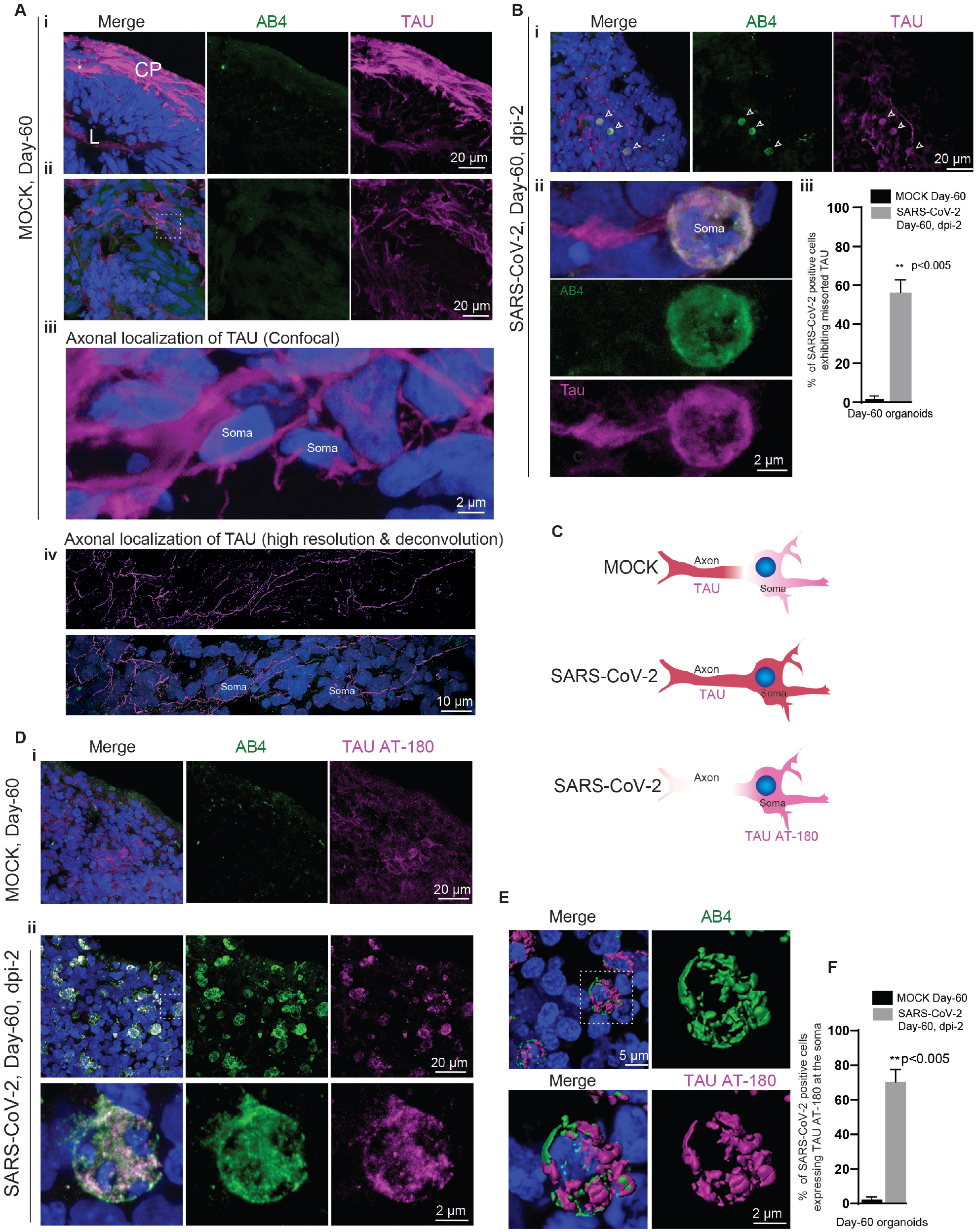
SARS-CoV-2 deregulates of Tau in cortical neurons. **A.** Tau immunoreactivity (magenta) specifies the cortical plate (CP) **(i).** Selected optic sections at high magnification **(ii)** and **(iii)** show Tau localization only in axons of cortical neurons. Note the somas of neurons are free from Tau protein. At least eight organoids from four different batches (n=4) are tested. Figures display scale bars. **B.** Tau missorting in SARS-CoV-2-positive neurons **(i).** Note, in contrast, to control groups, SARS-CoV-2-exposed organoids display missorted Tau (magenta) majorly into the somas of neurons (arrowheads). Selected confocal slices are shown to distinguish TAU missorting into neuronal soma. At high magnification, neuronal soma is further specified by the perinuclear localization of SARS-CoV-2 (green) **(ii).** Bar diagram at right **(iii)** quantifies the percentage of SARS-CoV-2-positive neurons exhibiting missorted TAU. At least six organoids from four different batches (n=4) are tested. Figures display scale bars. **F.** The bar diagram quantifies the fraction of missorted Tau neurons among with SARS-CoV-2-positive neurons. For statistics, at least 300 cells from four organoids and two independent batches (n=2) were examined. Unpaired t test with Welch’s correction, **P<0.005. Error bars represent mean + SD. **C.** Schematic cartoon of differential Tau distribution in mock compared to SARS-CoV-2-positive neurons. In mock, Tau is sorted mainly to axons. In SARS-CoV-2 positive neurons, Tau is missorted to the soma (determined by Pan-Tau antibody). Furthermore, phosphorylated Tau (at T231) majorly localizes in the soma (bottom panel, determined by TAU AT-180 antibody, see below). **D.** In contrast to controls **(i),** TAU AT180 antibody (magenta) that specifically recognizes the phosphorylated Threonine 231 of Tau protein distinctly localizes at the somas of SARS-CoV-2-positive neurons (green) **(ii).** At least four organoids from two different batches (n=2) are tested. Figures display scale bars. **E.** Co-localization of SARS-CoV-2 (green) and phosphorylated Tau protein (magenta) at somas of cortical neurons revealed by high-resolution imaging and de-convolution. Representative images from at least 300 cells examined. Figures display scale bars. **F.** The bar diagram quantifies the fraction of AT180-positive neurons co-localize with SARS-CoV-2-positive neurons. For statistics, at least 250 cells from four organoids and two independent batches (n=2) were examined. Unpaired t test with Welch’s correction, **P<0.005. Error bars represent mean + SD.

Compared to control organoids where Tau normally localizes in axons, SARS-CoV-2-positive neurons exhibited a strikingly different Tau localization pattern, although it was challenging to visualize Tau missorting in 3D tissues. Nevertheless, using selected confocal sections, we could image Tau missorting in SARS-CoV-2-positive neurons. In particular, we noticed an enhanced level of Tau missorted into the somas of the SARS-CoV-2-positive neurons **(Figure 3B-C).** To efficiently visualize Tau missorting, which is challenging in 3D tissues, we exposed iPSC-derived cortical neurons to SARS-CoV-2 in 2D. In contrast to unexposed cultures, SARS-CoV-2-positive neurons distinctly displayed Tau missorting such that most of the Tau localized to the soma of neurons **(Figure S3).**

During the pathogenesis of Alzheimer’s disease and other Tauopathies, Tau also gets hyper-phosphorylated at multiple sites. Sequential phosphorylation at different sites ultimately leads to hyperphosphorylation of Tau (*31*). Phosphorylation of Threonine 231 (T231) is the first event in the cascade of phosphorylation. This posttranslational modification specifically associates with non-fibrillar pre-neurofibrillary tangles found in the cytoplasm, somas, and nuclei of neurons in the brain tissue of Alzheimer’s disease (*33–35*). More precisely, compared to control organoids, early Tau-phosphorylation marker AT180 strongly localized at the soma of the SARS-CoV-2-positive neurons **(Figure 3D-F)**.

We then imaged cortical neurons for additional phosphorylated Tau using AT8 antibodies (specific for S202 and T205 of Tau) and p396 (specific for S396 of Tau). Although they label the axons of SARS-CoV-2-exposed organoids with slightly increased intensity, unlike AT180, none of these phospho-specific antibodies recognized Tau in the soma of SARS-CoV-2-positive neurons **(Figure S4).** In summary, these results demonstrate the presence of very specific phosphorylation of Tau at position T231 and the missorting of Tau to the soma of SARS-CoV-2-positive neurons implying neuronal stress reactions upon virus entry.

### SARS-CoV-2 induces neuronal cell death

Phosphorylation of Tau at T231 allows for isomerization of the following proline residue into distinct *cis-* and *trans-* conformations by the propyl-isomerase PIN1 (*36*). Cis-pT231Tau is acutely produced by neurons after traumatic brain injury, leading to disruption of the axonal microtubule network and apoptosis (*37*) (*38*). Accordingly, analyzing the nuclei of SARS-CoV-2-positive cells **(Figure 4A)**, we realized that they are highly condensed or fragmented exhibiting a strong reaction to 4’,6-diamidino-2-phenylindole (DAPI) that labels nuclei, a feature quite frequently observed in dead cells. To test neuronal cell death as a consequence of SARS-CoV-2 infection, we stained the SARS-CoV-2-exposed samples with Terminal deoxynucleotidyl transferase dUTP nick end labeling (TUNEL) that detects fragmented DNA in dead cells. We identified a significant number of cell death, suggesting that SARS-CoV-2-positive cells have undergone cell death within 2-dpi **(Figure 4B)**.

**Figure 4.**
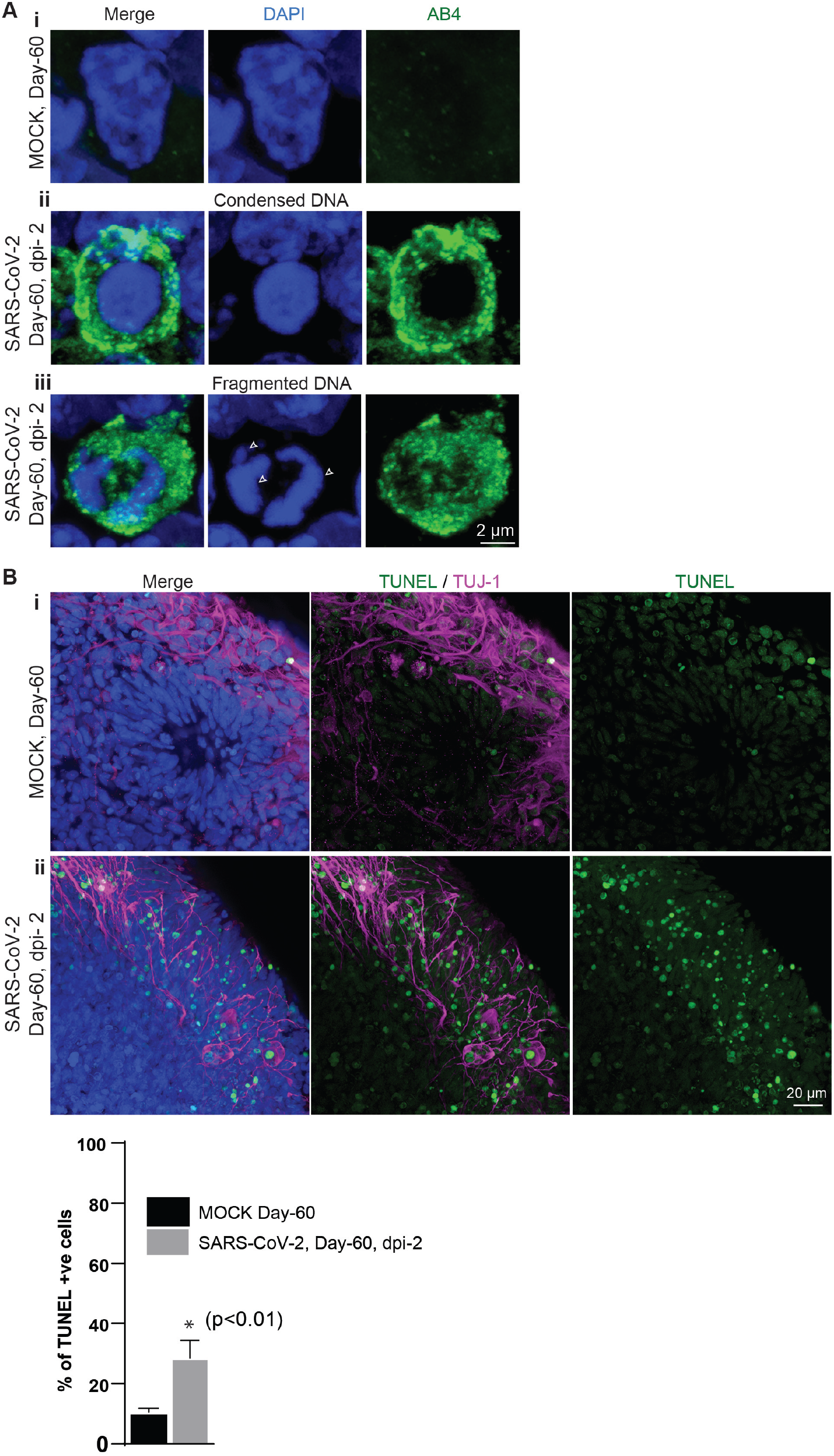
SARS-CoV-2 induces of neuronal death. **A.** Cells from mock-organoids display a healthy nucleus labeled by DAPI (blue) **(i).** SARS-CoV-2-positive cells (green) display condensed (middle panel, ii) and fragmented DNA (bottom panel, iii, arrows). At least 75 cells from two (n=2) independent batches of organoids were examined. Figures display scale bars. **B.** Compare to mock organoids **(i)** SARS-CoV-2-exposed organoids **(ii)** display increased TUNEL-positive cells (green) at the cortical plate that is specified by TUJ-1 (magenta). At least four organoids from two (n=2) independent batches of organoids were examined. Figures display scale bars. The bar diagram quantifies the frequencies of TUNEL-positive cells between mock and SARS-CoV-2-exposed organoids. Four organoids from two (n=2) independent batches were examined. Unpaired t test, *P<0.05. Error bars represent mean + SD.

## Discussion

We used 3D human brain organoids as an experimental system to study the COVID-19 and identified unexpected neurodegeneration-like effects of SARS-CoV-2 on human neurons. So far, the possible direct effect of SARS-CoV-2 on the CNS has been debated but not experimentally demonstrated. Thus, it was essential to examine whether SARS-CoV-2 can directly target human neurons and whether this leads to productive infection.

The finding that SARS-CoV-2 preferentially targets cortical neurons but not actively proliferating NPCs may suggest that developing embryonic brains are potentially less susceptible or free from the neurotoxic effects of SARS-CoV-2. This is indeed in striking contrast to ZIKV, a neurotropic virus that directly infects NPCs and triggers them to prematurely differentiate into neurons leading to congenital microcephaly (*20*) (*21*) (*22*).

In contrast to vascular, kidney, and intestinal organoids (*27–29*), SARS-CoV-2 cannot actively replicate in brain organoids at least until 4-dpi. There are several reasons why the human brain organoids might not support the active replication of SARS-CoV-2. Firstly, the developmental stages of brain organoids used in this work may not contain the full complement of SARS-CoV-2’s host cell replication factors. As an example, efficient replication of SARS-CoV requires ACE-2 (*39*) whose expression appears to be relatively low in brain organoids **(Figure 2G).** Next, the cell tropism of SARS-CoV-2 in brain organoids is post-mitotic neurons, which may not support the generation of viral progeny **(Figure 2B-C).** Future experiments using aged organoids and bioengineered organoids with SARS-CoV-2 replication factors are required to conclude if brain organoids can support productive infection of SARS-CoV-2.

ACE-2 is an entry receptor for SARS-CoV, and efficient replication of SARS-CoV (SARS outbreak in the year 2003, also depends on the expression level of ACE-2 (*39, 40*). Curiously, SARS-CoV could only infect the brain of transgenic mice expressing an elevated level of human ACE-2 but not non-transgenic mice. This key finding suggests that the neurotropism of SARS-CoV, to some extent, depend on the expression level of human ACE-2 in the brain (*12*). From this, although unexpectedly, the brain is susceptible to SARS-CoV-2 infections, our experiments using human brain organoids reveal that human neurons are indeed vulnerable to SARS-CoV-2 infections even though they express a low level of ACE2. This finding offers a couple of possibilities. First, even a basal level of ACE2 expression is sufficient for viral entry into the neurons. Second, the presence of yet unknown neuron-specific viral entry factors has to be elucidated. Even a low level of ACE-2 is sufficient for the viral entry is an intriguing phenomenon as it could explain why SARS-CoV-2 has a broad spectrum of target organs and cell types (*13*).

Our findings reveal that SARS-CoV-2 does not merely infect cortical neurons but has a consequence. SARS-CoV-2 infection induces pathological effects similar to early Tauopathies and neuronal cell death. Detection of early Tau phosphorylation at T231 in SARS-CoV-2-positive neurons is remarkable as it can trigger a cascade of downstream effects that finally could initiate neurodegenerative-like diseases. Early Tau phosphorylation could be reversible (*31*). However, phosphorylation events observed in conjunction with apparent neuronal cell death suggest that SARS-CoV-2 has detrimental effects on cortical neurons **(Figure 3 and 4).** Another interesting aspect to consider is that the current pandemic is the acute phase of SARS-CoV-2 infection. Thus, it is unclear what chronic or long-term effects it may cause in the CNS.

In conclusion, COVID-19 research has taken center stage in biomedical research. It is noteworthy that three coronavirus epidemics have occurred within the last two decades, and thus the future zoonotic coronavirus outbreak cannot be unexpected. With the advent of emerging human organoid research, which did not exist twenty years ago, we should be able to model the current SARS-CoV-2 infections and sufficiently prepare us for the future. Recent works utilizing kidney and gut organoids have already revealed insights into infection mechanisms (*27–29*). Adding to them is the current work that establishes brain organoids as a test system for SARS-CoV-2 infection and provides indications for potential neurotoxic effects of SARS-CoV-2. Since organoids are experimentally tractable human in vitro system and convenient to culture as well as to infect, organoid systems may well serve as a test-bed to screen for anti-SARS-CoV-2 agents. The presented work only provides initial insights. Future experiments utilizing mature state of brain organoids, bioengineered organoids, and orthogonal experiments with complementary *in vivo* experimental models are assured to dissect the neuropathology of SARS-CoV-2.

## Acknowledgment

We want to thank Dr. Boris Görg for offering generous support with their microscope facility. We want to thank Ms Gladiola Goranci and Nazlican Altnisk for their excellent technical assistance. This work was financially supported by a grant from Fritz-Thyssen Foundation. “We would like to thank the diagnostics department of the Institute of Virology, University Hospital Düsseldorf. Authors declare that they have no competing financial interests.

## List of supplementary materials

### Methods

#### Clinical specimens

For the isolation of infectious SARS-CoV-2 particles, nasopharyngeal, and oropharyngeal swab specimens from one individual with positive qRT-PCR results for SARS-CoV-2 infection was used. The Swab specimen was transported in a viral cultivation medium and stored at 4°C overnight. Freezing at −20°C was found to interfere with the infectivity of viral particles. Before the inoculation of susceptible cells, 500 μL maintenance medium (Dulbecco’s Modified Eagle Medium (Thermo Fisher), 2 % Fetal Calf Serum (PAN Biotech), 100 U/mL Penicillin and 100 μg/mL Streptomycin (Gibco) was added to the swab specimen. To get rid of major impurities, samples were briefly centrifuged (3000 x g; 60 sec) and the supernatant was transferred to new vials.

#### Human respiratory epithelial cells (hREC) and culturing

To obtain respiratory epithelia, a MedScand Cytobrush Plus GT (Cooper Surgical, Trumbull, USA) with a gentle-touch tip was rinsed with isotonic saline before use. Afterward the brush was inserted into the inferior nasal meatus followed by rotatory and linear motions against the medial and superior side. Isolated cells were transferred into a 15 mL centrifuge tube (Corning Incorporated, New York, USA) with 5 mL prewarmed RPMI 1640 medium containing 2 % Antibiotic-Antimycotic 100 x (Gibco^®^ Life Technology, Grand Island, USA). The brushes were vigorously shaken several times within the tube and cells were pelletized by centrifugation at 900 rpm for five minutes at room temperature. hRECs were re-suspended in Dulbecco’s Modified Eagle Medium/Nutrient Mixture F-12 (DMEM/F-12, Gibco^®^ Life Technology, New York, USA) supplemented with 2 % UltroserTM G Serum Substitute (Pall Corporation, Port Washington, USA) and 2 % Antibiotic-Antimycotic 100 x and seeded on T-25 or T-75 rat-tail collagen-coated tissue flasks (Greiner Bio-One, Kremsmünster, Austria), according to the pellet size respectively, and incubated at 37°C, 5 % CO2.

To reduce the risk of contamination, the medium was replaced after 24 hours, and the flasks were then integrated into the regular feeding procedure (exchange of medium every 48 – 72 hours). After one week, the concentration of Antibiotic-Antimycotic was reduced to 1 %. Reaching confluency of 90 %, the collagen layer was digested by incubating with 200 U/mL collagenase type IV (Worthington Biochemical Company, New Jersey, USA) for 30 – 60 min, followed by several washing steps with DMEM/F-12 supplemented with 1 % Antibiotic-Antimycotic. To reduce the number of fibroblasts, the pellet was re-suspended in 7 mL DMEM/F12 supplemented with 2 % UltroserTM G, seeded on tissue culture treated T-25 flasks (Corning Incorporated, New York, USA) and incubated for 1 hour at 37°C, 5 % CO2. The cells were then separated by incubating with Trypsin-EDTA 0,05 % for five minutes before the reaction was stopped with FBS followed by centrifugation at 900 rpm for five minutes at room temperature.

After re-suspending in PneumaCultTM-Ex Medium (STEMCELLTM Technologies, Vancouver, Canada), 4 x 105 cells/mL were seeded on collagen-coated 6,5 mm Transwell^®^, 0.4 μm pore Polyester membrane inserts (Corning Incorporated, New York, USA) with 250 μL medium on the apical side and 500 μL on the basolateral side, respectively. Before airlift, after three to five days, depending on cell confluency, PneumaCultTM-Ex Medium was replaced every day at the apical and basolateral side. To perform airlift, the medium on the apical side was carefully removed, whereas the basolateral medium was exchanged with PneumaCultTM-ALI Medium. The airlifted inserts were then integrated into the regular feeding procedure and incubated at 37°C, 5% CO2. Fully differentiated, the pseudostratified epithelium is expected 15-30 days after airlift and resembles human airway epithelium (in vivo) with respect to function and morphology.

#### Inoculation of Vero cells

In compliance with the decision of the German committee on biological agents (ABAS) of the Federal Institute for Occupational Safety and Health, all experimental studies involving infectious SARS-CoV-2 were performed within the biosafety level 3 (P3) facility at the University Hospital Düsseldorf. For isolation of SARS-CoV-2 from a clinical specimen, 2.5×105 Vero cells (ATCC-CCL-81, obtained from LGC Standards) were seed into T25 cell culture flasks in maintenance medium and cultured at 37°C in a humidified cell culture incubator. The following day, SARS-CoV-2 inoculum was prepared by diluting 200 μL of a clinical specimen with 800 μL maintenance medium. The medium was removed from Vero cells, and 1 mL inoculum (1 mL of maintenance medium for control Vero cells) was added onto the Vero cell monolayer. Vero cells were incubated for one h on a laboratory shaker at 37°C in a humidified incubator. Afterward, 4 mL of maintenance medium were added. To monitor viral replication, 100 μL of supernatant were directly harvested as the first sample (0 h post-inoculation) and every 24 h for four days post-inoculation. Additionally, cells were imaged by light microscopy.

#### Real-time qPCR analysis for quantification of SARS-CoV-2 RNA copies per mL

For extraction, 100 μL cell culture supernatant was incubated with 400 μL AVL buffer (viral lysis buffer used for purifying viral nucleic acids; cat No. 19073, Qiagen, Hilden Germany) for 10 min at RT and mixed with 400 μL 100 % ethanol. RNA extraction was performed with 200 μL cell culture mix using the EZ1 Virus Mini Kit v2. (cat. no. 955134, Qiagen, Hilden, Germany) following the manufacturer’s instructions. A total of 60 μL were eluted from the 200 μL starting material. Five μL of the eluate were tested in qRT-PCR using the real-time TaqMan^®^-technique. A 113-base pair amplicon in the E-gene of SARS-Cov-2 was amplified and detected, as described by Corman et al (*41*) with minor modifications. The thermal protocol described has been shortened to 40 cycles of 95° C. We used the LightMix^®^ Modular SARS and Wuhan CoV E-gene (Cat.-No. 53-0776-96) and the LightMix^®^ Modular EAV RNA Extraction Control. We used the AgPath-ID^®^ One-Step RT-PCR Kit (Applied Biosystems, Cat. No. 4387391). RT-PCR was performed with an ABI 7500 FAST sequence detector system (PE Applied Biosystems, Weiterstadt, Germany). As a DNA-standard, a plasmid (pEX-A128-nCoV2019-E-gene) that encompasses the amplified region was created and serially diluted after purification. The software constructed a standard graph of the CT values obtained from serial dilutions of the standard, the CT values of the unknown samples are plotted on the standard curves, and the number of SARS-CoV-2 RNA copies was calculated.

For gene expression analysis of ACE2, quantitative RT-PCR analysis was performed by using qPCR MasterMix (PrimerDesign Ltd) and fluorescence emission was monitored by LightCycler 1.5 (Roche). For normalization, primers #5163 (5’ CCA CTC CTC CAC CTT TGA 3’) and #5164 (5’ ACC CTG TTG CTG TAG CCA 3’) were used monitoring cellular GAPDH expression. Expression was then calculated as 2^(-Δct)^.

#### Propagation of infectious SARS-CoV-2 particles

For propagation of infectious SARS-CoV-2 particles from Vero cell culture supernatant, 2.5×105 Vero cells were seeded into T25 cell culture flasks in maintenance medium and incubated at 37°C in a humidified cell culture incubator. The next day, the supernatant of inoculated Vero cells at day four post-inoculation (see above) was diluted with maintenance medium (1:2, 1:10, 1:100, 1:1000) in a total volume of 5 mL and added to the cells, which were incubated for four days at 37°C.

#### Determining SARS-CoV-2 viral titer by TCID50 assay in 96-well plates with Vero cells

For determination of viral titer in TCID50/mL, 5*10^3^ Vero cells were seeded in the first ten columns of 96-well plate in 100 μL maintenance medium and incubated at 37°C in a humidified cell culture incubator for 24 h. In a new 96-well plate, 180 μL maintenance medium was added to all wells of the first ten columns. For serial dilutions of the virus stock, 20 μL of the stock solution were added to the wells of the first column. Then 20 μL of the first dilution were transferred to the wells of the next column to obtain ten-fold serial dilutions up to 10^-9^. The tenth column of the 96-well plate serves as a control. After exchanging the medium of the previously prepared Vero cell plate with 100 μL fresh maintenance medium, 100 μL of each virus dilution were transferred to the Vero cell plate. After incubation at 37°C for four days, microscopic inspection of the plate was used to monitor cytopathic effects (CPEs) in the form of detached cells. TCID50/mL was determined as:

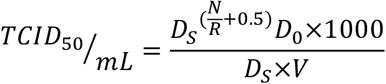

D_S_ = Dilution factor of consecutive dilutions (10); N = Total number of wells showing CPE; R = Replicates per dilution (8); D_0_ = Dilution factor of the first dilution (10); V = Volume per well in μL (200 μL).

#### SARS-CoV-2 infection

All experiments including SARS-CoV-2 infections were performed in aP3 safety laboratory (see above). Number of viral particles used to infect cells or organoids is identical for all three strains. Neurons and brain organoids were tested and found free from mycoplasma contamination. For infection with SARS-CoV-2, 25- and 60-day old organoids were transferred from spinner flasks into low-adherent 12 well plates. Each well contained one organoid in 2 mL differentiation medium and infected with SARS-CoV-2. was incubated as stationary suspension culture and treated. To exclude the effects observed were not induced by SARS-CoV-2, the control organoids (control, uninfected) were treated with supernatants of non-infected Vero cells.

#### Generation of convalescent serum and ELISA validation

AB1 and AB2 were obtained 23 and 16 days after the diagnosis of SARS-CoV-2 infection. AB3 and AB4 were obtained 27 and 28 days after the diagnosis of SARS-CoV-2 infection (by PCR). 1. Blood samples were drawn directly into serum collection tubes and spun for 15 minutes at 3.000 rpm. After centrifugation, the clear supernatant was aliquoted and stored at −80°C. ELISA was performed using semi-quantitative SARS-CoV-2-IgA and SARS-CoV-2-IgG ELISAs that detect binding against the recombinant S1 domain of the SARS-CoV-2 spike protein (Euroimmun, Lübeck, Germany).

#### Generation of iPSCs-derived cortical neurons

We differentiated iPSCs into NPCs using STEMdiff Neural Induction Medium (Stem cell technologies, USA). Five days later, the formed neurospheres were collected and cultured on poly-L-ornithine (PLO)-/laminin coated dishes. Seven days later, using a neural rosette selection medium (Stem cell technologies, USA), we re-cultured neural rosettes to generate NPCs. NPCs were differentiated into cortical neurons as described previously (*37*). Briefly, NPCs were seeded on poly-L-ornithine (PLO)-/laminin coated coverslips. Forty-eight hours later, NPCs were switched to cortical neuronal differentiation medium consisting of BrainPhys basal medium(*38*) supplemented with 1x B27 (without vitamin A, Thermo scientific, USA), 1x N2 (Thermo scientific, USA), 20ng/mL BDNF (Peprotech, USA), 20ng/mL GDNF (Peprotech, USA), 20ng/mL NT3, 1μM cAMP (Sigma, USA) and 0.2μM ascorbic acid (Sigma, USA). Fresh medium was added every 2-3 days.

#### Generation of iPSCs-derived brain organoids

Organoids were generated from two different iPS cell lines, namely IMR90 (Donor 1) and Crx iPS (Donor 2) as described previously (*42*). We adapted previously described protocol to differetiate iPSCs into brain organoids described earlier (*15, 17*). Five days old neurospheres were harvested and embedded in matrigel (Corning, USA) drops. Differentiation medium mixture of DMEM/F12 and Neural Basal Medium (in 1:1 ratio), supplemented with 1:200 N2, 1:100 L-glutamine, 1:100 B27 w/o vitamin A, 100 U/mL penicillin, 100 μg/mL streptomycin, 23 μM insulin (Sigma-Aldrich), 0.05 mM MEM non-essential amino acids (NAA), and 0.05 mM β-mercaptoethanol (Life Technologies) was used to differentiate the matrigel embedded droplets in suspension culture. After four days of culturing, embedded neurospheres were transferred to spinner flasks (IBS, Integra biosciences) containing the same differentiation medium supplemented with 0.5 μmol dorsomorphin (Sigma-Aldrich, USA).

#### Western blot

The gel electrophoretic separation of proteins was performed under denaturing conditions in the presence of SDS in a non-continuous gel system, which consisted of a 5% stacking gel and 10% resolving gel, which was then transferred to nitrocellulose membranes. Once the transfer was finished, the membrane was soaked into 5% milk in TRIS/HCl based buffer (TBST) for a minimum of 30 min at RT. After incubating with primary antibodies overnight at 4°C, the blots were treated with secondary antibodies at RT for 1 h. Super Signal West Pico or Femto Chemiluminescent substrates (Pierce) were used for detection. Anti-body dilutions for Western blots: Human convalescent serum AB4 (1:400), polyclonal rabbit anti SARS COV-2 (1:500), monoclonal mouse anti SARS COV-2 (1:500), and mouse anti GAPDH (1:20,000, 60004-I-Ig, Proteintech). As the secondary antibodies goat anti-Mouse IgG (H+ L) HRP (1:5000, 31430, Thermo Fisher Scientific), goat anti-Rabbit IgG (H+L) HRP (1:5000, 31466, Invitrogen), Anti-human secondary antibodies conjugated to HRP (1:5000, Thermoscientific).

#### Immunofluorescence and confocal microscopy

For light microscopy analysis, monolayer cells (Vero and aspics-derived neurons) were fixed for 10 minutes. Brain organoids were fixed for 30 minutes. We used 4% paraformaldehyde/PBS as a fixative (*17*). Organoids were incubated in 30% sucrose overnight at 4°C, embedded in Tissue-Tek O.C.T. compound (Sakura, Netherlands). Organoids were cryofrozen at −80°C before sectioning into 10 to 15 μm thin slices using Cryostat Leica CM3050 S. Thin sections and cells were permeabilized with a buffer containing 0.5% Triton X100 for 10 minutes. Specimens were blocked with 0.5% fish gelatin/PBS for 1 hr, both at room temperature. For SOX2 stainings, antigen retrieval was required. For this, sections were treated with repeated heating (microwave) in Sodium citrate buffer (10 mM Sodium citrate, 0.05% Tween 20, pH 6.0) was applied before permeabilization and blocking.

We used different antibodies as follows: Human convalescent serum (AB1 to AB3 (1:40), AB4, 1:40 or 1:50 or 1:100), rabbit anti-TUJ-1 (1:400, Sigma Aldrich, USA), AT180 (1:100, MN1040, monoclonal mouse anti-phospho-Tau, AT8 (1:100, Thermo Fisher), polyclonal rabbit anti-phospho-Tau (S396), (1:100, Thermo Fisher), monoclonal mouse anti-SARS-CoV-2 (1:200, GeneTex), rabbit anti-SARS-CoV-2 NP (1:200, Biozol), rabbit anti-MAP2 (1:100, Proteintech), mouse anti-Pan-Tau (5A6, 1:100, DSHB). Specimens with primary antibodies were incubated overnight at 4°C. For secondary antibodies, Alexa Fluor Dyes conjugated either with goat/donkey anti-mouse, antihuman, or anti-rabbit (1:500 or 1:1000, molecular probes, Invitrogen) was used. For DNA staining, DAPI at a concentration of 1 μg/mL (Sigma Aldrich, USA) was used, and the coverslips were mounted using Mowiol (Carl Roth, Germany). The raw images were collected using a Leica SP8 confocal system (Leica microsystems, Germany) and processed with the help of Adobe Photoshop (Adobe Systems, USA). For deconvolution, the captured image files were processed using ZEN software (2.3, SP1, black, 64bit, release version 14.0.0.0; ZEISS, Oberkochen, Germany) for 3D reconstruction and deconvolution. After deconvolution, files were imported into Fiji and further processed using Image J, Adobe photoshop CC 2018, and Adobe Illustrator CC 2018. For 3D surface and volume rendering, raw image files were processed using Imaris (x64 version 7.7.1).

#### TUNEL assay

Apoptotic cells were detected by using DeadEnd™ Fluorometric TUNEL System (Promega, G3250, USA) according to the manufacturer’s protocol.

#### Ethical approval and patient samples

Serum samples AB1 and AB2 were obtained under a protocol approved by the ethical commitee, medical faculty, University Hospital Düsseldorf, Heinrich-Heine-University (study number 5350). Serum samples AB3 and AB4 were obtained under a protocol approved by the Institutional Review Board of the University of Cologne (protocol 16-054). Human respiratory epithelial cells (hREC) were obtained by nasal brush biopsy from healthy control individuals. The study was endorsed by the local ethical committee at the University of Münster, and each patient gave informed written consent (Study number, 2015-104-f-S, Flimmerepithel) and 2020-274-f-S (COVID-19). Trained physicians from the Department of General Pediatrics, University Hospital of Münster, performed biopsies.

### Supplementary figure legends

**Figure S1.**
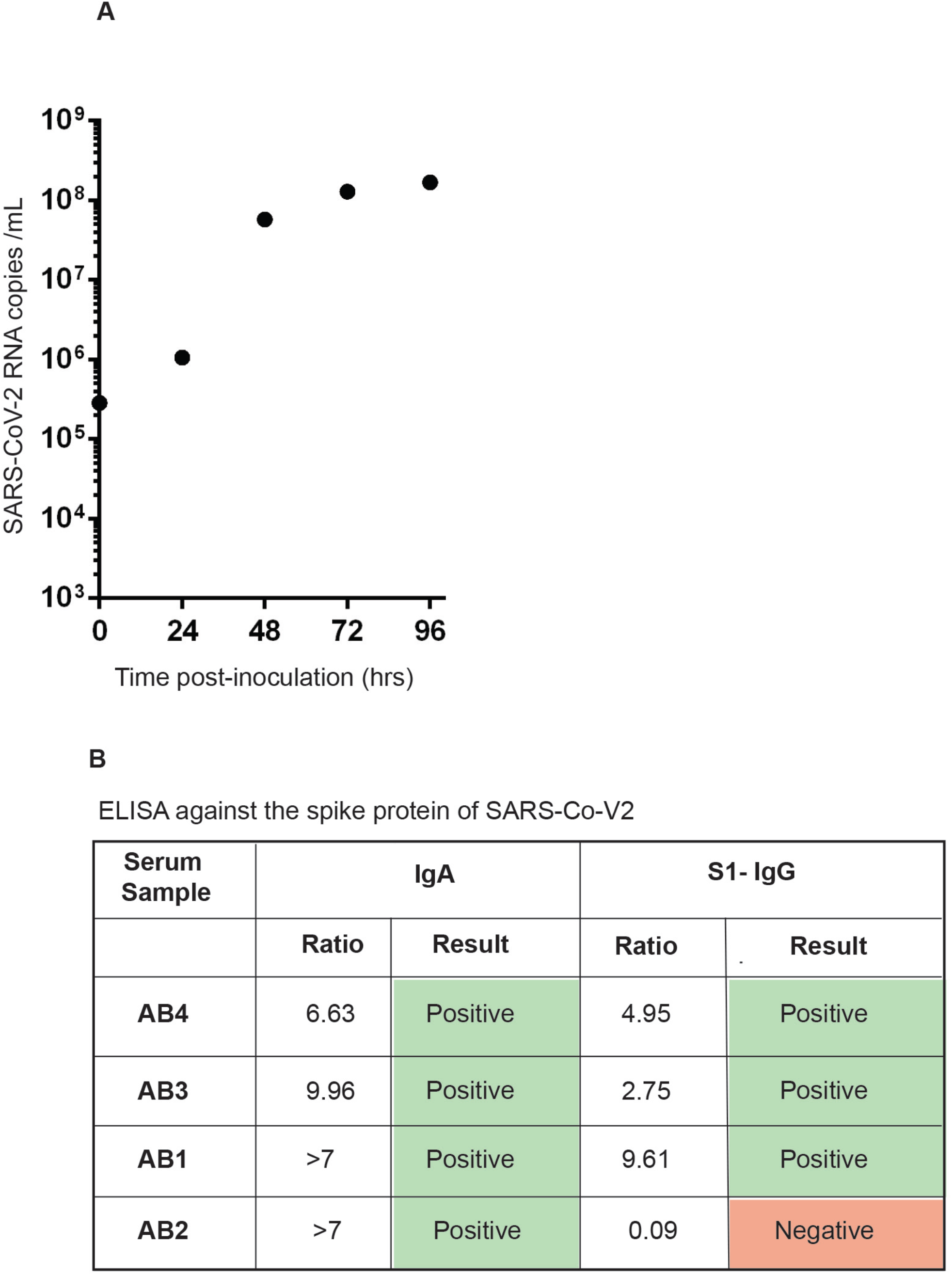
Assaying of SARS-CoV-2 replication and convalescent serum (Related to Figure 1) **A.** SARS-CoV-2 productively replicates in inoculated Vero cells. Real-time quantitative polymerase chain reaction (qPCR) analysis of Vero cell culture supernatant shows that SARS-CoV-2 RNA drastically increases from 0-dpi until 3-dpi. **B.** Isolation of COVID-19 convalescent serum and their analysis in an enzyme-linked immunosorbent assay (ELISA). Except for AB2, the rest of the convalescent serums contained IgG.

**Figure S2.**
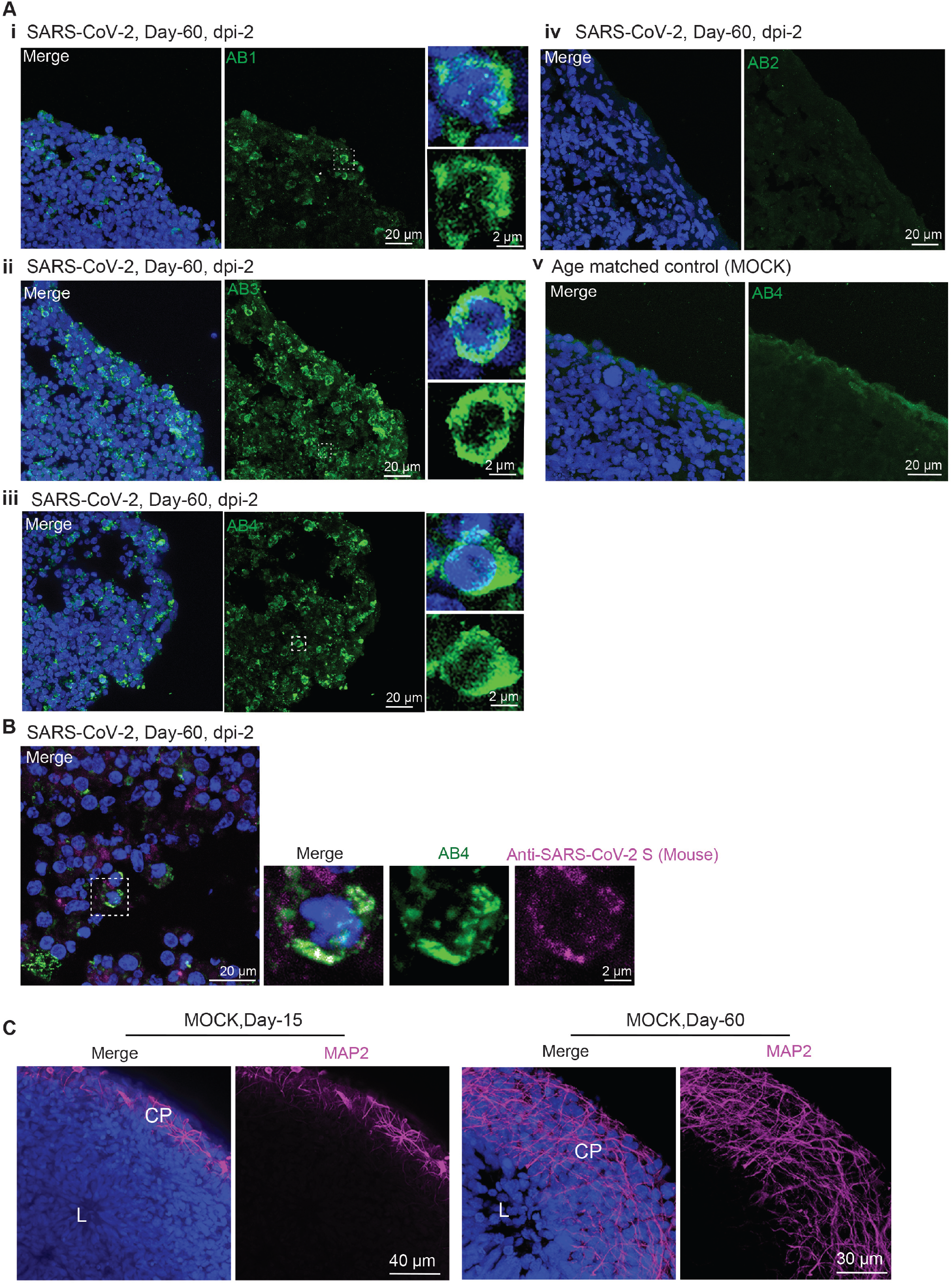
The convalescent serum AB4 detects SARS-CoV-2-positive cells. **A.** Immunostaining detection of SARS-CoV-2 in organoids exposed to SARS-CoV-2. Serum AB1, AB3, and AB4 detect sub-population of cells (green). In contrast, AB1 does not recognize cells in SARS-CoV-2-exposed organoids. None of these serums identify cells in control un-exposed organoids. At least three independent organoids were examined. Figures display scale bars. **B.** Further validation of AB4. Mouse monoclonal Anti-SARS-CoV-2 S detects (magenta) SARS-CoV-2-positive cells labeled by AB4 (green). Figures display scale bars. **C.** Compared to Day-15 organoids **(i),** Day-60 organoids **(ii)** exhibit signs of cortical maturation as distinguished by the abundance of MAP2-positive neurons in their cortical plate (CP). Note an increased size and thickness of MAP2-positive cortical plate in Day-60 organoids.

**Figure S3.**
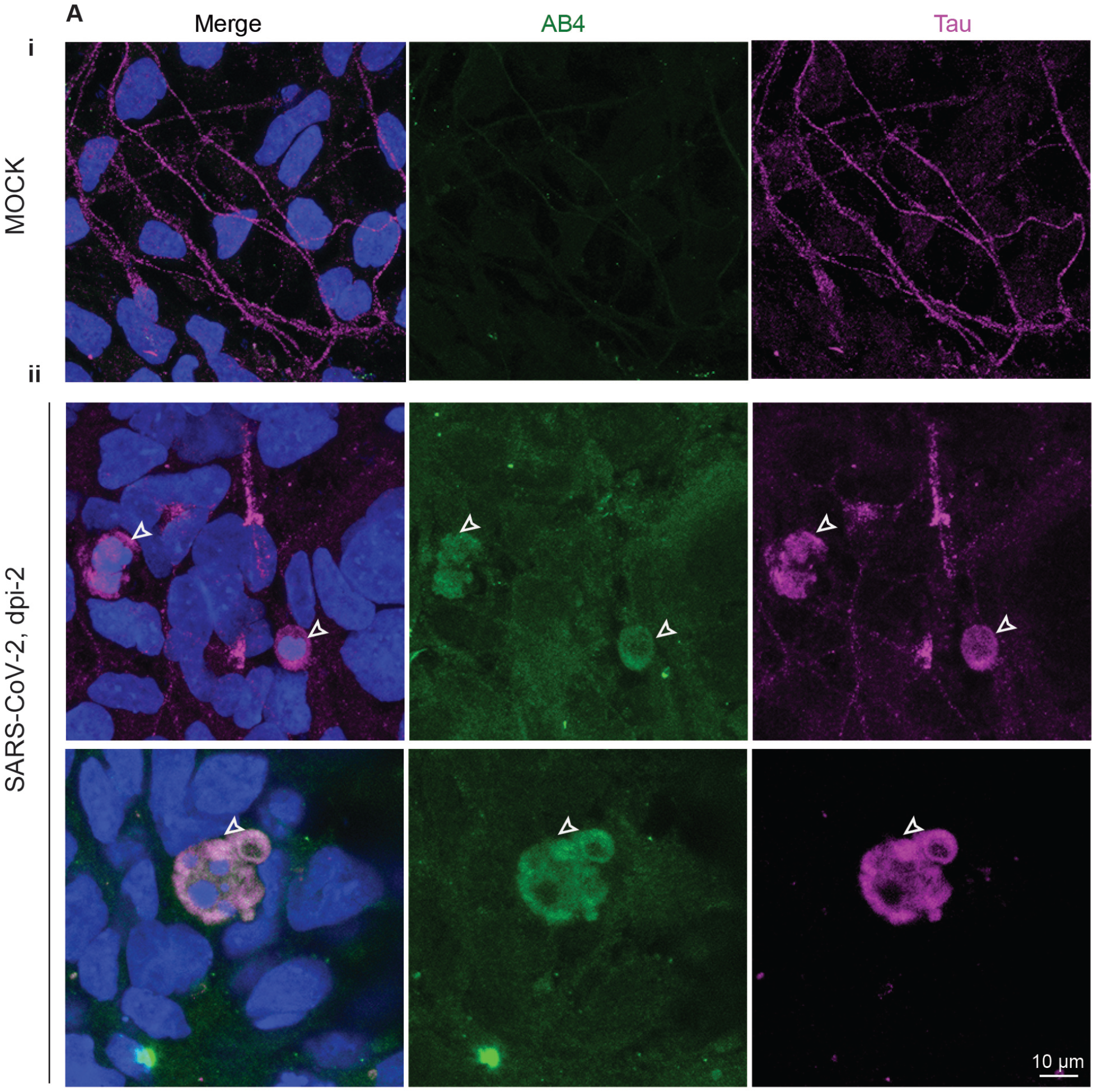
Tau missorting in 2D experiments (Related to main Figure 3) **(A)** Tau immunoreactivity (magenta) specifies the axons of iPSC-derived cortical neurons in 2D **(i).** In SARS-CoV-2-exposed 2D cultures (**ii),** SARS-CoV-2 (green) localizes at the soma of neurons (magenta, arrowheads). In this condition, Tau mostly mislocalized to cell soma. At least 50 neurons were examined from two independent (n=2) experiments. Figures display scale bars.

**Figure S4.**
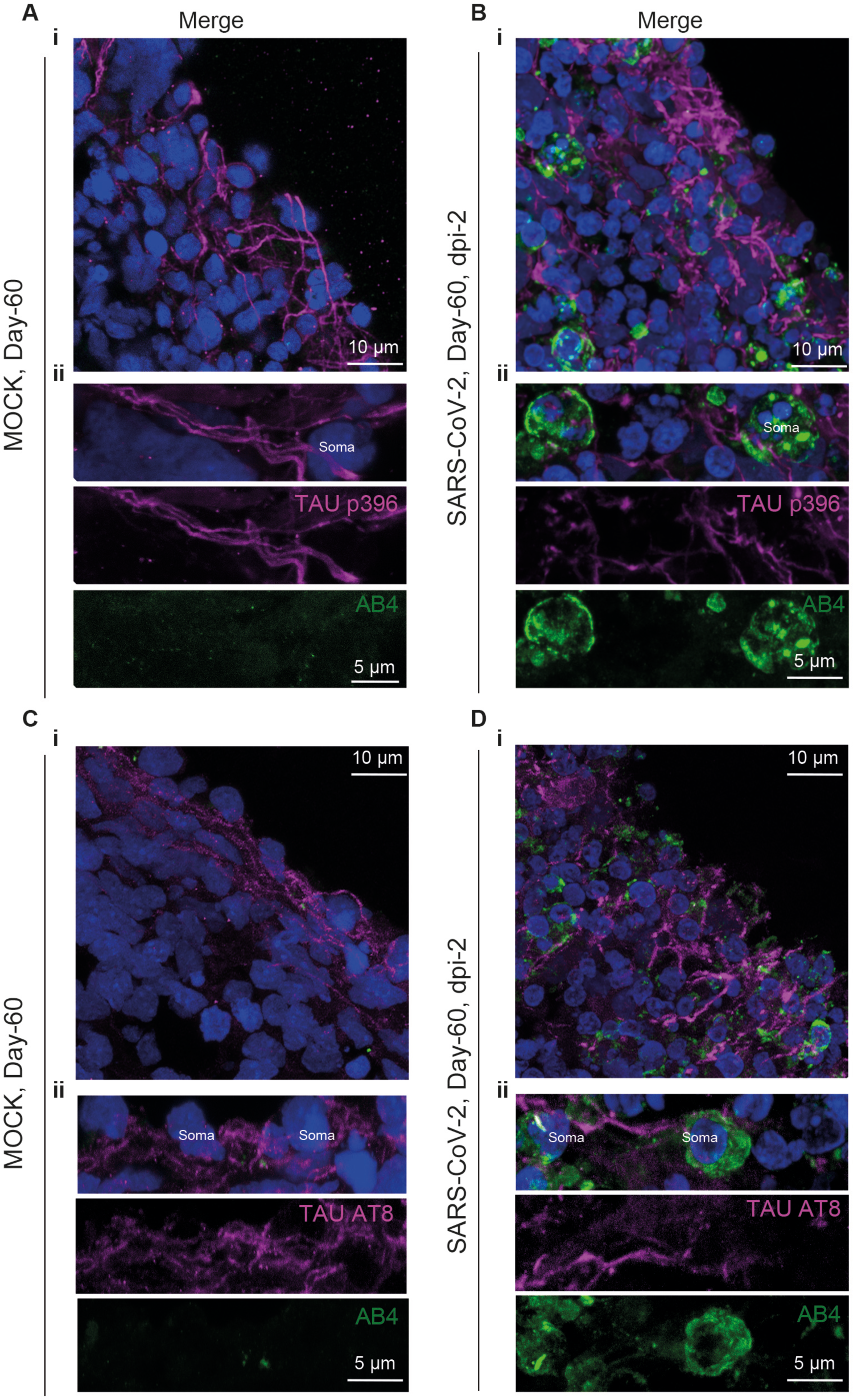
Tau S396 and Tau S202/T205 do not localize into somas of SARS-CoV-2-positive neurons. **A.** Tau p396 antibody that labels phosphorylated Tau at S396 localizes to axons of Day-60 (magenta, mock organoids) and do not localize to soma of SARS-CoV-2-positive neurons (green) **(B).** At least four organoids from two different batches (n=2) are tested. Figures display scale bars. **C.** AT8 antibodies that labels phosphorylated Tau at S202 and T205 localizes to axons of Day-60 (magenta, mock organoids). This antibody does not recognize Tau in soma of SARS-CoV-2-positive neurons (green) **(D).** At least four organoids from two different batches (n=2) are tested. Figures display scale bars.

